# Computational Prioritization of T Cell Epitopes to Overcome HLA Restriction and Antigenic Diversity in *Plasmodium falciparum*

**DOI:** 10.1101/2025.07.14.664425

**Authors:** Alexander J. Laurenson, Brian G. Pierce, Shannon Takala-Harrison, Matthew B. Laurens

**Affiliations:** Center for Vaccine Development and Global Health, University of Maryland School of Medicine, Baltimore, Maryland, USA; Molecular Microbiology and Immunology Program, Graduate Program in Life Sciences, University of Maryland School of Medicine, Baltimore, Maryland, USA; University of Maryland Institute for Bioscience and Biotechnology Research, Rockville, Maryland, USA; Department of Cell Biology and Molecular Genetics, University of Maryland, College Park, Maryland, USA

**Keywords:** *Plasmodium falciparum*, malaria, epitope prediction, immunoinformatics, multi-epitope vaccine

## Abstract

Developing a highly effective malaria vaccine remains challenging due to *Plasmodium falciparum*’s antigenic diversity and human leukocyte antigen (HLA) polymorphisms, which complicate vaccine antigen selection and limit immune protection. The first recommended malaria vaccine, RTS,S, provides only partial, allele-specific protection with waning immunity over time, and the more recently developed R21 vaccine will likely encounter the same hurdles. To address these challenges, we developed a computational tool that integrates *P. falciparum* sequence diversity, predicted T cell epitope-HLA binding affinities, and HLA allele frequencies from sub-Saharan Africa to identify conserved, immunogenic epitopes with broad population coverage. We analyzed 42 *P. falciparum* proteins, previously identified as vaccine candidate antigens, and generated consensus sequences using data from 18 African countries, and then incorporated HLA allele frequencies from 24 sub-Saharan populations. CD8+ and CD4+ T cell epitopes were predicted using NetMHCpan-4.1 and NetMHCIIpan-4.1. Our novel tool, T cell Epitope Nomination (TEpiNom), used greedy optimization to filter and select epitopes based on epitope sequence conservation (>95%), binding affinity (median rank <10%), and broad HLA coverage, minimizing redundancy to reduce immune escape risk. Our tool identified 2,265 MHC I and 1,992 MHC II conserved epitopes spanning pre-erythrocytic, erythrocytic, and sexual stage proteins. Key MHC I epitopes from pre-erythrocytic antigens HSP70-2, SLARP/SAP1, p36, FabZ, LISP1, LSA1, UIS3, p24_2, PL, and FabG achieved near 100% HLA-A, HLA-B, and HLA-C coverage, and MHC II epitopes from pre-erythrocytic, erythrocytic, or sexual antigens provided 98.5%-100% coverage for a given parasite life stage. This strategy advances malaria vaccine design by integrating epitope promiscuity and multistage antigen selection to support broad, durable protection and identify promising multi-epitope malaria vaccine candidates for subsequent experimental validation. Our computational framework is adaptable for vaccine development against other genetically diverse and immunologically evasive pathogens.

## Introduction

The global malaria death toll declined from 897,000 in 2000 to 577,000 in 2015, but has remained stagnant since then, with an estimated 597,000 malaria deaths in 2023, 76% of which occurred in children less than 5 years of age [1]. This plateau occurred despite the implementation of multiple public health interventions, underscoring the urgent need for improved malaria control measures, including more effective vaccines.

The very first malaria vaccines, RTS,S/AS01 and R21/Matrix-M, were recently recommended by the World Health Organization [2]. RTS,S and R21 target the Circumsporozoite Protein (CSP) present on circulating sporozoites during the pre-erythrocytic stage of infection [3]. While these vaccines have demonstrated moderate efficacy, extended Phase III trials revealed at least three limitations: 1) RTS,S provides allele-specific immunity, 2) efficacy wanes over time without booster doses, and 3) protection is lower in younger children [3]. Long-term protection for the R21 vaccine remains uncertain [4]. Despite the significant public health benefit of these vaccines, the need for more effective vaccines remains elusive for multiple reasons. One contributing factor includes the parasite’s extensive genetic diversity, which is not comprehensively represented by a monovalent vaccine and can result in allele-specific immune responses [5,6]. Another reason that remains understudied is HLA-restricted vaccine efficacy, which can make the successful development of vaccine-mediated immunity between individuals with different HLA alleles widely variable [7].

Malaria vaccines are categorized by the parasite life stage they target. Pre-erythrocytic vaccines, which include the RTS,S and R21, can target the parasite in its sporozoite form during the migration to or invasion of hepatocytes after human injection by a mosquito during a blood meal [3,4]. This class of vaccines can also target the parasite during development and replication within hepatocytes before blood stage (erythrocytic) infection. Pre-erythrocytic vaccines are highly favorable as inhibition of parasite invasion and development at this stage avoids infection entirely. Infected hepatocytes express antigens that alert CD8+ T cells to kill infected cells, while infected erythrocytes cannot, as they lack the endogenous antigen-presenting MHC I protein. This capacity to manipulate the cytotoxic arm of cell-mediated immunity makes the pre-erythrocytic stage even more attractive as a vaccine target [6].

Erythrocytic vaccines target malaria parasites after they are released from the liver as merozoites into the bloodstream to infect erythrocytes [8,9]. These vaccines aim to stimulate an immune response that reduces the number of parasitized red blood cells, including targeting merozoite surface proteins essential to red blood cell invasion. By limiting the quantity of infected red blood cells, erythrocytic vaccines reduce the clinical severity of infection and assist in clearing parasites [7]. Transmission-blocking vaccines target parasites after their differentiation into sexual forms by blocking either gamete fertilization or zygote development into sporozoites in the mosquito vector [6]. Although people immunized with a transmission-blocking vaccine can experience infection and clinical disease, the vaccine should still One promising strategy that could successfully halt parasite transmission either from human to mosquito or from mosquito to the next person during a blood meal [6].

One major challenge in malaria vaccine development is identifying suitable gene products and proteins to target, given the size and complexity of the *Plasmodium falciparum* (*Pf*) genome. With almost 5,300 genes, many of which encode proteins involved in immune evasion and host invasion, pinpointing the most conserved and immunologically relevant antigens is a daunting task [10,11]. Since the first *Pf* reference genome was assembled in 2002, characterization of putative functions of gene products has advanced, and genetic diversity amongst circulating strains has been extensively described [10,12,13]. The MalariaGEN Pf7 database is a curation of over 16,000 quality-controlled *Pf* genomes from 33 countries, including more than 8,000 from sub-Saharan Africa, where ∼95% of malaria fatalities occur [1,14]. This extensive amount of genomic data has transformed the malaria vaccine research landscape, enabling a more precise investigation of candidate antigens that are both conserved and immunogenic across diverse populations.

Conventional vaccines that utilize live or attenuated pathogens can require years to develop, mostly spent fine-tuning the balance of inactivating a pathogen without limiting its protective immunogenic potential [6]. Subunit vaccines, including RTS,S and R21, often rely on a single haplotype sequence of a single protein to stimulate protective immune responses. As demonstrated for candidate erythrocytic vaccines and for RTS,S, when a vaccinated individual encounters a parasite strain with a dissimilar sequence in the target protein, that parasite may escape the vaccine-induced immune response [5,6,15–18]. Similar to how widespread antimalarial drug pressure has repeatedly selected for drug-resistant parasite variants, circulating *Pf* strains could evade allele-specific vaccine-induced immunity if only a single protein domain is targeted [8]. Given these limitations, novel malaria vaccine approaches are needed to achieve broader and more durable protection.

One promising strategy that could overcome the limitations of conventional vaccine design, especially in the context of malaria, is reverse vaccinology, an approach that begins with the pathogen’s genome to systematically identify vaccine candidate antigens [19]. Pioneered by Dr. Rino Rappuoli, this method revolutionized vaccine development by enabling antigen target down-selection informed by genetic information without initial experimental screening [19]. Reverse vaccinology is particularly appropriate for addressing the unique hurdles presented by complex pathogens such as *Pf*. An extension of reverse vaccinology is the design of epitope-based vaccines, which focus on the inclusion of short peptide sequences derived from pathogenic proteins that elicit robust T or B cell immune responses.

Epitope-based vaccines are particularly well-suited for targeting genetically diverse pathogens because they enable inclusion of multiple highly conserved epitopes, regions that remain stable across strains and may therefore be less likely to mutate under immune pressure. These epitopes can be screened en masse based on preferential attributes such as sequence conservation and binding promiscuity to HLA alleles (for T cell epitopes) or surface accessibility (B cell epitopes). Promising epitope candidates characterized as highly conserved and putatively immunogenic could then be experimentally validated before inclusion in tandem in an epitope-based vaccine. By discretely targeting dominant epitopes from separate, nonredundant protein antigens, epitope-based vaccines offer a strategic advantage for eliciting broad and durable immunity against complex and antigenically variable pathogens such as *Pf* [20].

Epitope-based malaria vaccines with higher efficacy and immunogenicity than existing vaccines can significantly enhance public health benefits, potentially requiring fewer doses, eliminating the need for annual boosters, and blocking onward transmission. Such advancements would boost individual protection, reduce the burden on vaccine delivery systems, improve cost-effectiveness, diminish transmission, and advance health equity. By carefully selecting epitopes and pairing them with effective adjuvants, epitope-based vaccines can be designed to elicit robust cell-mediated and humoral immune responses that protect against infection [21].

In the context of malaria, the activation of key T cell subsets is critical for effective immunity, yet it remains understudied relative to B cell activation and antibody responses. CD8+ T cells play a role in eliminating intracellular parasites by targeting and killing infected hepatocytes before a *Pf* parasite enters the bloodstream (Figure 1A) [22,23]. The cytotoxic response that targets infected hepatocytes is essential for pre-erythrocytic immunity, as infected hepatocytes represent a ‘bottleneck’ stage during which parasites are especially vulnerable to immune clearance. Meanwhile, CD4+ T cells are required to coordinate immune responses by differentiating into T helper (Th) cell subsets, such as IFNγ-producing Th1 cells, which enhance macrophage function and amplify innate immune defenses, effector functions considered integral to clearing intracellular parasites during liver and erythrocytic stages (Figure 1B) [24,25]. CD4+ Th1 T cells also contribute to refining the humoral response by producing cofactors that induce B cell class switching [26]. Additionally, another subset of CD4+ T cells, T follicular helper (Tfh) cells, provide essential co-stimulatory signals that drive B cell activation within germinal centers, promoting development of high-affinity, long-lived plasma cells and memory B cells, and ensuring sustained adaptive immunity and effective parasite clearance (Figure 1C) [27,28]. Given the essential role of T cells in malaria immunity, especially at the pre-erythrocytic stage, designing vaccines that effectively stimulate both CD8+ and CD4+ responses is a critical step toward enhancing protection and blocking transmission.

**Figure 1:**
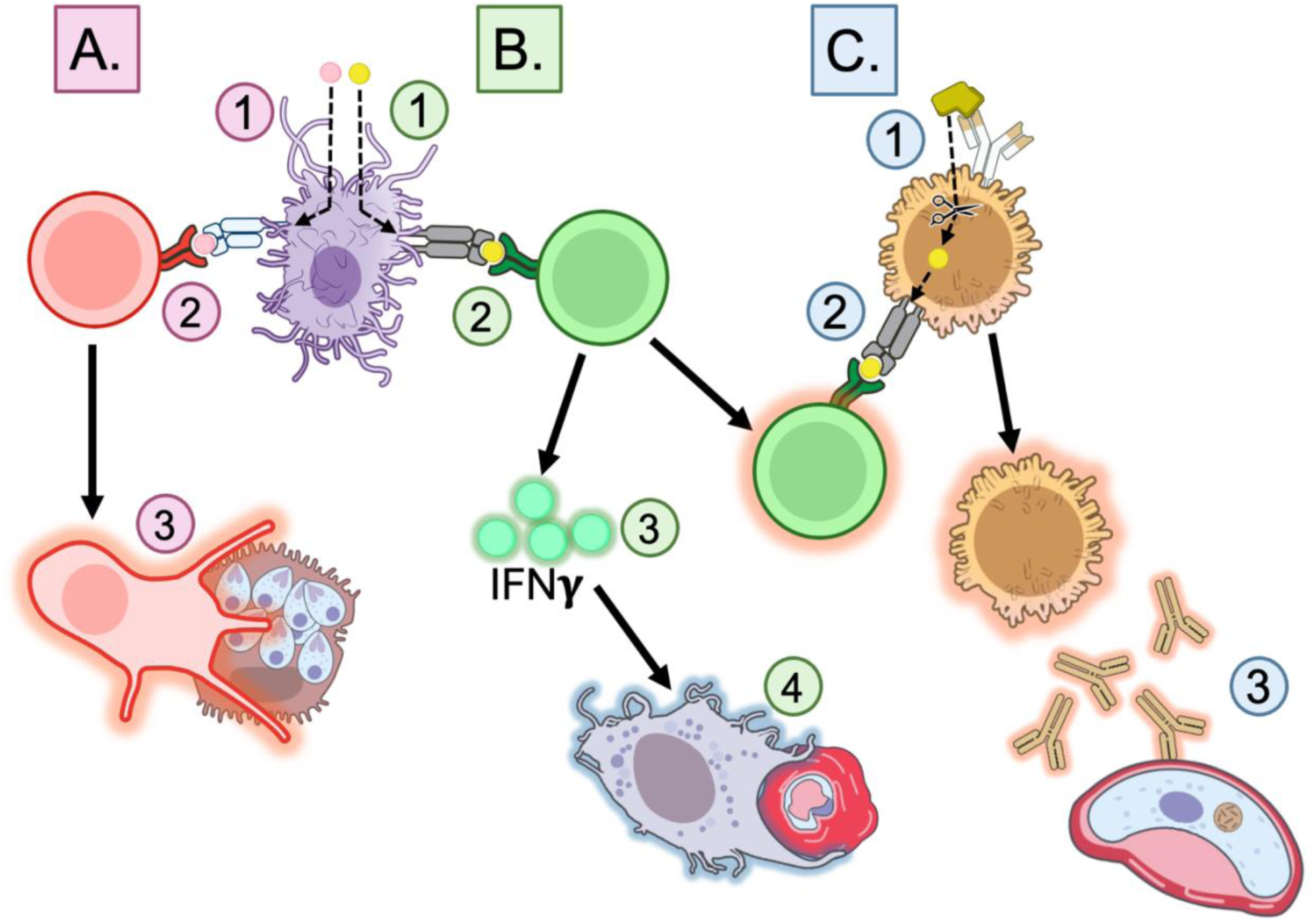
Epitope-induced immunity. A. CD8+ Cytotoxic T cell activation. 1. MHC I epitope uptake and processing by Antigen-Presenting Cell (APC). 2. Epitope cross-presentation on MHC I protein and binding to CD8+ T cell receptor. 3. Killing of sporozoite-infected hepatocytes by CD8+ Cytotoxic T cell. **B. CD4+ Helper T cell activation.** 1. MHC II epitope uptake and processing by APC. 2. Epitope presentation on MHC II protein and binding to CD4+ T cell receptor. 3. Production of IFNγ by CD4+ Helper T cell. 4. Killing of merozoite-infected red blood cells by IFNγ-activated innate immune cells. **C. B cell activation.** 1. B cell epitope binding to B cell receptor and antigen uptake. 2. Antigen processing and presentation of epitope on MHC II protein. 3. Activated CD4+ Helper T cell binds to epitope on MHC II and co-stimulates B cell. 4. Activated B cell produces antibodies specific to sexual stage protein. Art is licensed under Public Domain and available on NIH BIOART.

To engineer an efficacious vaccine, T cell epitopes must first be selected based on their ability to bind with high affinity to Major Histocompatibility Complex (MHC) I and MHC II, the proteins responsible for presenting peptides, short amino acid sequences that comprise epitopes, to T cells. MHCs in humans are encoded by HLA (Human Leukocyte Antigen) genetic loci, which are exceedingly diverse, specifically within the region that encodes the peptide-binding groove [29,30]. This high degree of HLA sequence diversity is particularly important when deciding which T cell epitopes to target with vaccines, as HLA allele frequencies differ significantly across populations. An epitope that binds strongly to HLA alleles common in one regional population may exhibit poor binding to common alleles in another population, leading to geographic disparities in vaccine efficacy. To address this challenge, it is critical to incorporate regional HLA polymorphism data when selecting vaccine epitopes, ensuring broad coverage across diverse populations. Since epitope binding is linear, computational sequence-based models can often accurately predict high-affinity epitope-HLA interactions, enabling the use of immunoinformatics for large-scale screening [20,31].

Recent work has demonstrated the feasibility and utility of integrating *in silico* predictions with experimental validation to down-select *Pf* T cell epitopes [32]. A study by Kotraiah et al. comprehensively analyzed five erythrocytic antigens for HLA-DRB1-restricted CD4+ T cell epitopes using epitope prediction tools and validated these restricted epitopes using *in vitro* binding assays and *ex vivo* recall response assays [32]. This work confirms the predictive accuracy of immunoinformatic approaches as an initial screen to identify specific, immunogenic T cell epitopes as putative vaccine targets, and illustrates the value of combining computational tools with experimental screening to yield actionable candidate epitopes for *Pf* vaccine design.

Despite the availability of parasite sequence data and HLA allele frequency data sets from endemic regions, these *in silico* approaches remain underutilized on a larger scale. A major barrier to implementing T cell epitope prediction tools is that they generate data sets with large volume outputs, requiring users to either manually filter data or rely on multiple separate tools for epitope down-selection, which makes the process arduous and often inaccessible. This study describes a systematic, immunoinformatics-driven approach to down-select predicted epitopes for *Pf* malaria vaccine development. Using parasite protein sequences from field isolates and HLA allele frequency data from endemic African populations, we apply computational prediction tools to identify T cell epitopes with strong HLA binding affinities within malaria proteins previously identified as essential to parasite survival or invasion (Figure 2). We then introduce a down-selection framework that filters these epitopes based on conservation, binding affinity, and HLA promiscuity, enabling prioritization of a manageable subset of highly immunogenic targets for experimental validation. Additionally, we incorporate a population coverage optimization step to select epitope combinations that maximize immune responses across diverse HLA alleles while minimizing protein redundancy in target epitope selection, reducing immune escape risk. This approach offers a systematic method for refining large-scale epitope predictions into a focused selection of high-priority candidates for *Pf* vaccine development.

**Figure 2:**
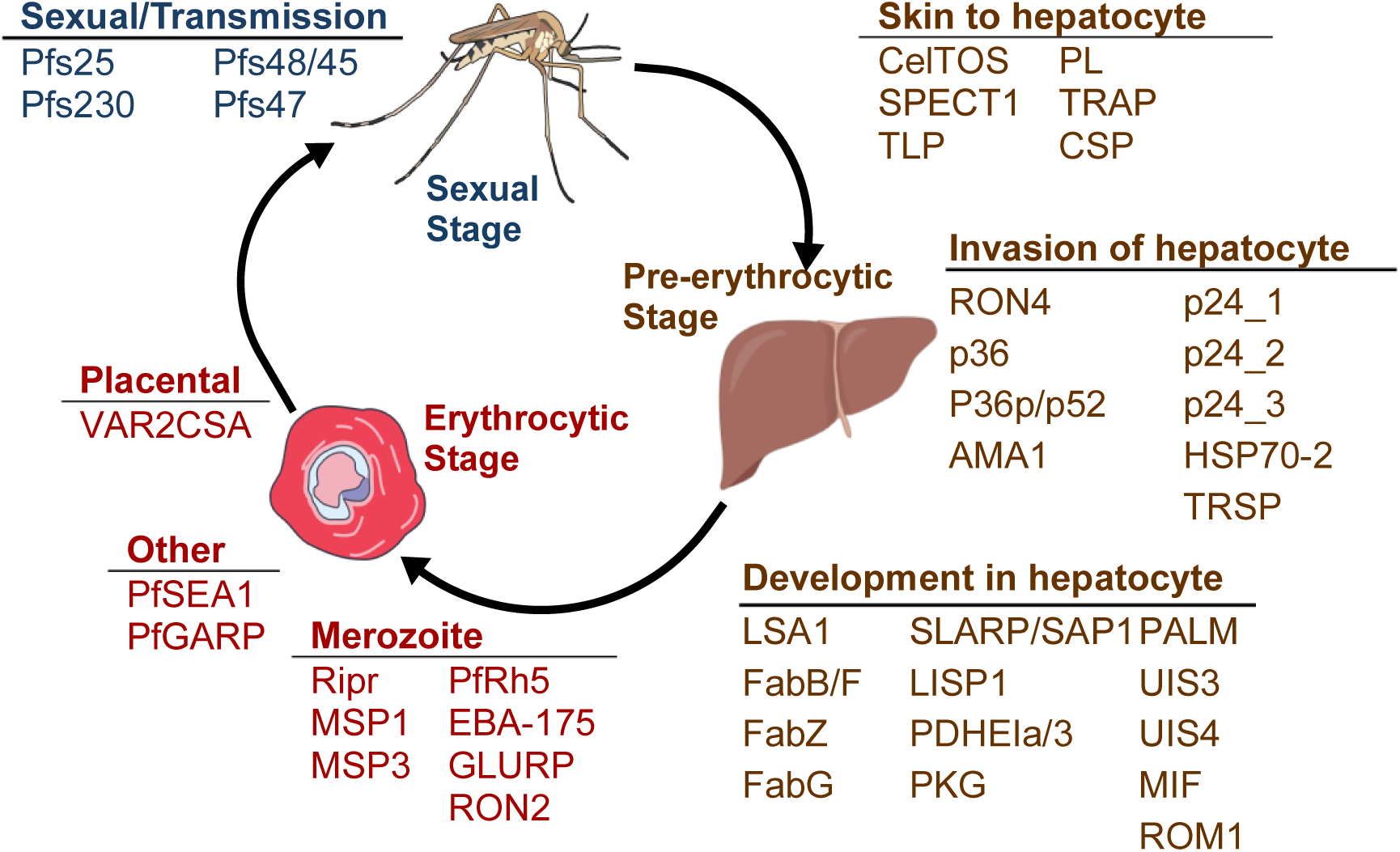
Vaccine candidates by malaria life stage. Liver, blood, and sexual/transmission stages of *Pf* and the vaccine candidates separated by stage and substage. Art is licensed under Public Domain and available on NIH BIOART and Bioicons.

## Methods

### Selection of Pf antigens and retrieval of sequences

Forty-two candidate proteins were selected based on prior research demonstrating their essential roles in *Pf* pathogenicity, particularly in parasite invasion or survival (Figure 2) [33,34]. Variant Call Files (VCFs), which had been mapped to the Pf3D7 reference genome, were acquired from publicly available MalariaGEN Pf7 and contained variants for *Pf* samples collected from malaria-experienced individuals. Samples within the VCFs that were collected within sub-Saharan African countries (Figure 3) and that were likely monoclonal infections, as indicated by a within-host infection fixation index (F_WS_) ≥0.95, were selected for consensus sequence generation [14]. After normalizing to the Pf3D7 reference genome, filtering adjacent insertions and deletions within 5 base pairs as per samtools and bcftools best practices, sample-specific variants were incorporated into the Pf3D7 reference sequence backbone to generate a full-length consensus sequence for each sample [35]. This process was repeated for each antigen. Sequences were translated using UCSC Genome Browser’s faTrans tool, and sample sequences with nonsense mutations were excluded [36]. The finalized protein sequence data sets, one for each of the 42 vaccine candidates, comprised information from 18 African countries with a median of 3,996 sample sequences per protein (Supplementary Table 2, Figure 3).

**Figure 3.**
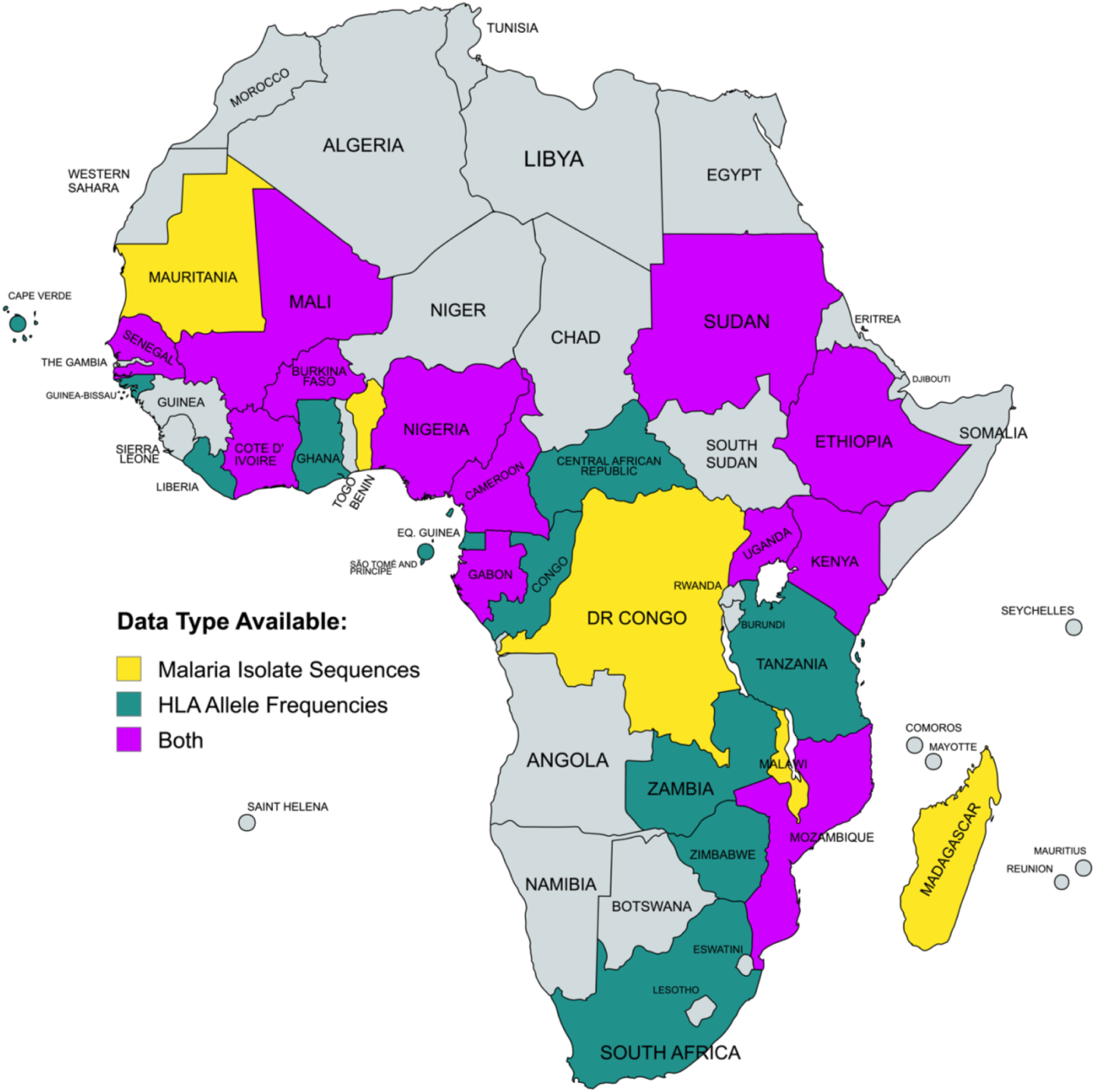
Parasite genomic and HLA data availability. Map of sub-Saharan African countries with MalariaGEN parasite data (red), HLA allele data (blue), or both data types (purple). Created with MapChart.

### Selection of HLA alleles and population frequencies

HLA allele frequency data was obtained by searching the Allele Frequency Net Database, a global repository for published MHC allele population frequency data, for HLA-A, HLA-B, HLA-C, HLA-DQ, HLA-DP, and HLA-DR alleles recorded within sub-Saharan African countries (Figure 3) [37–39]. Extracted allele frequencies, representing the proportion of each allele within each population sample, were normalized within each HLA class to reflect the relative frequencies of HLA alleles. The finalized HLA allele data set comprised information from all 24 countries with published HLA data sets, encompassing frequencies for 748 unique HLA alleles (Figure 3).

### Diversity analysis of protein antigens

Multiple Sequence Alignments (MSA) were performed within each protein candidate sequence database using MAFFT (v7.427) with default parameters [40]. After alignment, sequences were clustered using CD-HIT (v4.8.1) at a 100% identity threshold to define unique haplotypes [41]. Haplotype diversity (*Hd*) was then calculated for each protein using the standard formula, 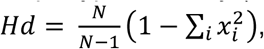 where *N* is the number of sequences and *x_i_* is the frequency of each haplotype in the sample.

### Prediction of T cell epitopes

T cell epitope binding to MHC I and II proteins was predicted using downloaded software for NetMHCpan-4.1 and NetMHCIIpan-4.1, neural network tools trained on both mass spectrometry binding affinity and eluted ligand data [31]. Epitope predictions were performed against the HLA-A, HLA-B, HLA-C, HLA-DQ, HLA-DP, and HLA-DR alleles within the previously mentioned HLA allele data set. CD8+ and CD4+ T cell epitopes were predicted as 8-11 and 15 amino acid long peptides, respectively. CD8+ T cell epitopes were predicted uniquely for pre-erythrocytic stage antigens and not for blood- or sexual stage antigens due to the lack of MHC I proteins on red blood cells.

### Design of T cell epitope down-selection tool

To streamline the down-selection of T cell epitopes, we developed a computational tool, T cell Epitope Nomination Tool (TEpiNom; Patent Pending) that integrates parasite protein sequence data sets, predicted epitope-HLA binding data, and HLA allele frequency data sets to down-select for or nominate a list of candidate T cell epitopes (Figure 4). Using Python and packages, pandas and NumPy, the tool filters and then ranks epitopes based on several immunological and practical criteria, including the predicted binding strength to HLA alleles, epitope sequence conservation across pathogen haplotypes, and HLA allele frequencies in human populations intended to be covered by the vaccine [42–44].

**Figure 4.**
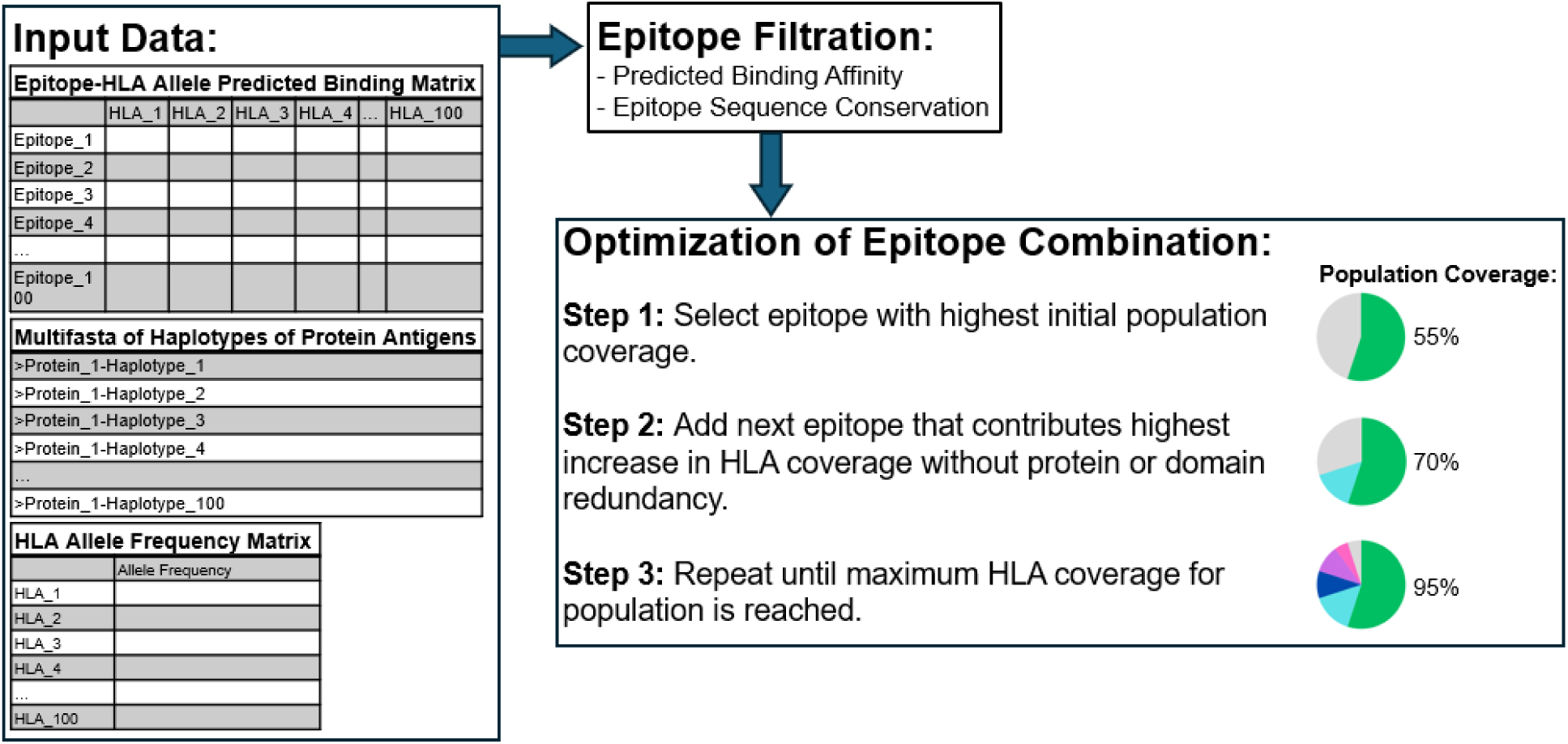
Workflow of T Cell Epitope Nomination (TEpiNom) algorithm. Input data includes epitope-HLA allele predicted binding matrix, multifasta of sequence for each protein antigen haplotype, and HLA allele frequency matrix. After epitope filtration using user-defined thresholds for median predicted binding affinity and epitope sequence conservation, the tool optimizes a combination of epitopes to maximize HLA coverage in the population, while minimizing target redundancy.

After filtration for epitope-HLA binding strength and epitope sequence conservation based on user-defined thresholds, the tool employs a greedy optimization algorithm to build optimal sets of epitopes that maximize HLA allele coverage while minimizing protein and domain redundancy. The algorithm considers an epitope predicted to bind to a given HLA allele within a user-defined threshold for binding affinity as coverage for that HLA allele by that epitope, therefore covering a certain percentage of the human population based on that HLA allele’s frequency in the population. The algorithm selects a starting epitope that allows for the highest initial population coverage based on the HLA alleles it’s predicted to bind to, and then iteratively adds epitopes that contribute the greatest marginal gain to HLA coverage, while also satisfying additional constraints on protein and domain redundancy. For example, once an epitope is selected from a particular protein or domain, other epitopes from the same protein or with overlapping sequences are deprioritized, reducing the intersection of immune pressure on the same protein or similar protein regions and improving antigenic breadth. The tool’s final output includes an epitope data set filtered based on specified conservation and rank binding affinity parameters, optimized epitope combinations with population coverage metrics for each HLA genetic locus, overall MHC I or II coverage, and circulating strain coverage based on epitope conservation.

### Filtration of Pf T cell epitopes

Using TEpiNom, the predicted MHC I and II epitopes within the malaria vaccine candidate proteins were filtered based on two main criteria: epitope sequence conservation and median predicted binding affinity ranks. The epitope sequence conservation (*C*) was calculated using the proportion of the protein sequence data set in which an epitope is 100% conserved, specifically with the following formula: 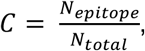 where *N_epitope_* is the number of sample sequences for the respective protein that contain the exact epitope sequence and *N*_*total*_ is the total number of sample sequences for that protein. Epitope binding affinity ranks are derived from NetMHCpan-4.0 and NetMHCIIpan-4.1, reflecting the predicted strength of epitope binding to specific HLA alleles (lower percent ranks indicate stronger predicted binding). The tool applied default defined cutoffs, retaining epitopes with epitope sequence conservation greater than 95% and median binding affinity ranks below 10% for further analysis. This filtration process, powered by the tool, resulted in a set of epitopes with high conservation and strong predicted binding affinities across the analyzed proteins, ready for population coverage optimization.

### Optimization of Pf T cell epitope combinations

To optimize the population coverage of T cell epitopes, TEpiNom was applied to identify the most promising combinations of epitopes for MHC I or II epitopes, with MHC II epitopes analyzed separately by protein life stage. First, epitope sequences and their predicted HLA binding affinities were compiled into a single data set. Using the tool, the population coverage of each predicted epitope was calculated by summing the percent population of sub-Saharan African individuals who have at least one HLA allele predicted to bind to a given epitope sequence, based on the generated HLA allele frequency data set. An epitope with a predicted binding affinity rank below 2% for an HLA allele was deemed to provide coverage for that allele. The tool then iteratively optimized combinations of epitopes, selecting up to ten epitopes per combination, to ensure that the final epitope set collectively maximized population coverage while prioritizing epitope selection from different proteins and protein domains.

### Analysis of Pf T cell epitopes by clinically associated HLA types

To investigate associations between HLA-restricted T cell epitopes and clinical outcomes, we conducted a systematic literature search to identify HLA alleles associated with key clinical outcomes: parasitemia, uncomplicated malaria, or severe malaria, including cerebral malaria. Searches were performed using PubMed with combinations of the following keywords: “*Plasmodium falciparum*,” “malaria,” “HLA,” “parasitemia,” “uncomplicated malaria,” “severe malaria,” and “cerebral malaria.” Inclusion criteria required studies to report HLA-allele associations with statistically significant outcomes in human cohorts, with data stratified by severity of disease. HLA data was extracted and compiled into a reference data set of HLA alleles classified as associated with a distinct clinical outcome. Predicted T cell epitopes for malaria vaccine candidates were analyzed for their binding affinity ranks to the HLA alleles identified in the literature for the prioritization of epitopes that bind promiscuously to protective and non-protective HLA alleles.

## Results

### Sequence diversity within candidate antigens

Haplotype diversity analysis across the protein sequence data sets (Supplementary Table 1) revealed varying levels of sequence diversity within each protein antigen (Figure 5). In the pre-erythrocytic stage, several proteins such as TRAP (0.997), AMA1 (0.996), CSP (0.981), and CelTOS (0.9799) displayed high haplotype diversity values, indicating substantial sequence variation amongst circulating strains (Figure 5a). Conversely, multiple pre-erythrocytic stage proteins, including HSP-70 (0.038), TRSP (0.052), PKG (0.055), and ROM1 (0.027), exhibited low haplotype diversity, suggesting a high degree of sequence conservation. Erythrocytic stage proteins also spanned a wide range of diversity, with MSP1 (0.996), GLURP (0.981), and EBA-157 (0.988) among the most diverse, while PfRh5 (0.617) and PfSEA1 (0.876) showed moderate diversity (Figure 5b). In the sexual stage, Pfs230 (0.999) was highly diverse, whereas Pfs25 (0.027) and Pfs48/45 (0.654) were more conserved (Figure 5c). These results demonstrate that while many candidate proteins are highly polymorphic, others show strong conservation, supporting the inclusion of conserved antigens and epitopes in vaccine formulations.

**Figure 5:**
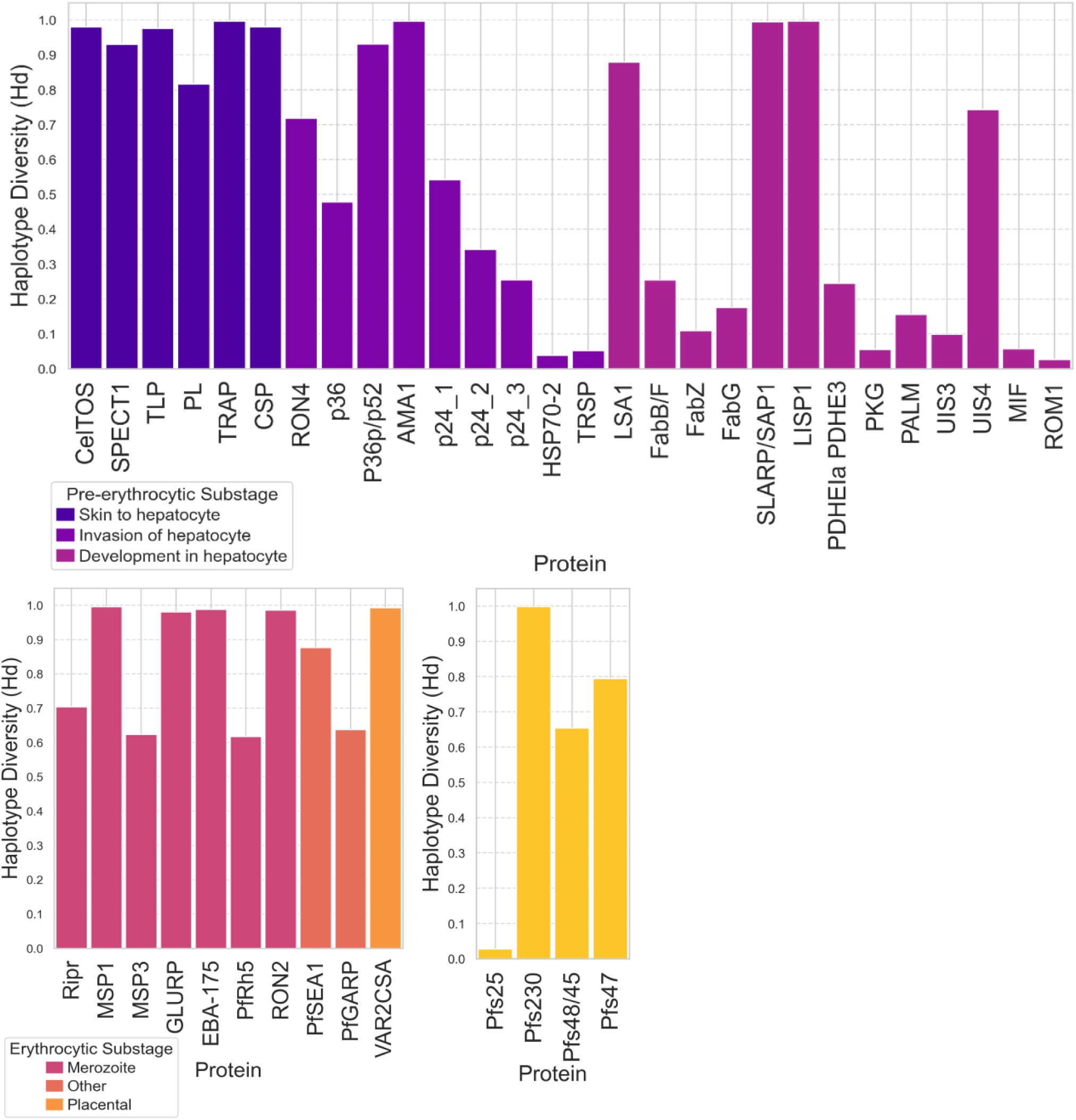
Haplotype diversity for malaria vaccine candidate proteins across different life cycle stages. Box plots show the distribution of haplotype diversity values for protein candidates within the **a. Pre-erythrocytic stage**, **b. Erythrocytic stage**, and **c. Sexual stage**, with sub-stages of protein function labeled. Haplotype diversity ranges from 0.00 to 1.00 and reflects the probability that two randomly chosen haplotypes from the population are different. Higher values indicate greater sequence diversity, while values closer to 0.00 reflect higher sequence conservation.

### Prediction and filtration of T cell epitopes within candidate proteins

To identify T cell epitopes within the protein antigens, we used the NetMHCpan-4.1 and NetMHCIIpan-4.0 tools to predict epitopes and their binding affinity ranks to MHC I and II proteins. Through this method, we predicted 244,036 unique MHC I and 164,034 unique MHC II epitopes, all with varying predicted binding affinity ranks to the input HLA alleles. We then used the TEpiNom workflow to filter epitopes based on their median binding affinity rank (<10%) to the HLA alleles and epitope sequence conservation (>95%). Predicted epitopes were retained only if they met both criteria.

After applying these filter thresholds, we identified 2,265 MHC I and 1,992 MHC II candidate epitopes across malaria vaccine candidate proteins and characterized the retention rate per protein, or the proportion of candidate epitopes that remained after filtration (Supplementary Table 4).

For MHC I epitope predictions, conducted exclusively for pre-erythrocytic antigens, multiple highly conserved and promiscuous epitopes were identified across all candidates (Figure 6a; Supplementary Table 2). ROM1 had the highest MHC I epitope retention rate (3.08%), followed by p36 (2.37%) and PALM (1.80%), suggesting these proteins contain relatively conserved, immunogenic regions. SLARP/SAP1 and LISP1 had the highest absolute numbers of retained MHC I epitopes, with 584 and 270 epitopes, respectively, indicating their strong potential as T cell targets.

**Figure 6:**
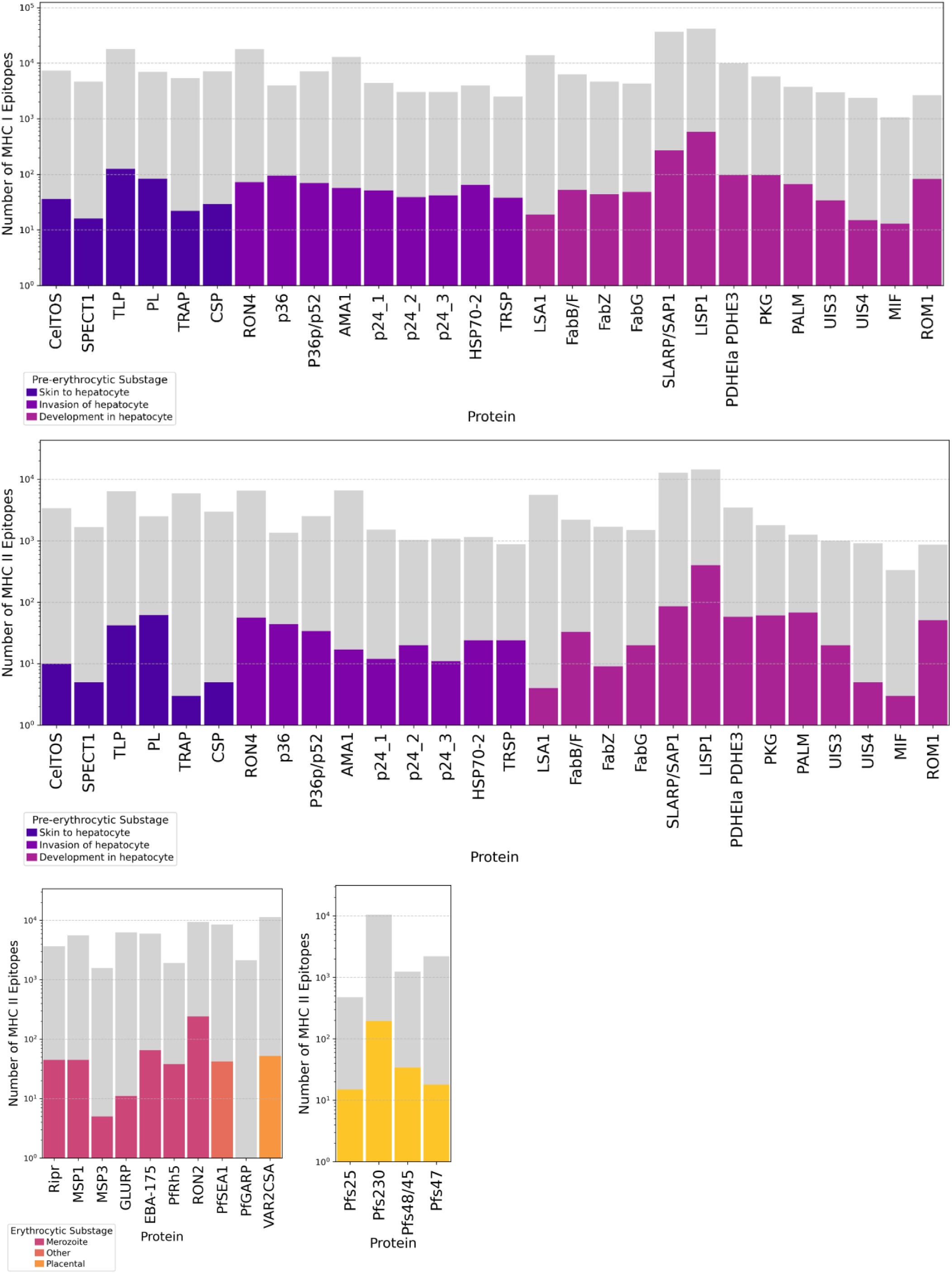
Predicted and retained T cell epitopes across malaria vaccine candidate proteins. Number of predicted MHC I and MHC II epitopes before and after applying conservation and binding affinity filters (<10% median binding rank, >95% conservation). Data is plotted on a logarithmic scale to show resolution and categorized by malaria life cycle stage (**a. Pre-erythrocytic MHC I epitopes, b. Pre-erythrocytic MHC II epitopes, c. Erythrocytic MHC II epitopes, and d. Sexual stage MHC II epitopes**) and substage with epitope counts shown for each protein after filtration steps. MHC I epitope predictions performed exclusively for pre-erythrocytic stage proteins.

For MHC II epitopes, which are critical for eliciting CD4+ T cell responses, the number and percentage of retained epitopes were similar to epitopes predicted to bind MHC I. Among pre-erythrocytic stage proteins, ROM1 (5.88%), PALM (5.40%), PKG (3.39%), and p36 (3.26%) had the highest retention percentages, suggesting their potential as strong CD4+ T cell inducers (Figure 6b; Supplementary Table 2). The pre-erythrocytic stage LISP1 had the highest absolute number of retained MHC II epitopes (400), followed by SLARP/SAP1 (86). Within erythrocytic antigens, RON2 retained the most MHC II epitopes (241, 2.56%), followed by PfRh5 (1.98%) and EBA-175 (1.09%) (Figure 6c; Supplementary Table 2). In contrast, GLURP, MSP3, and PfGARP had extremely low retention rates (<0.5%), indicating either greater sequence variability or weaker predicted binding affinities. For sexual stage antigens, retention rates were highest for Pfs25 (3.15%), Pfs48/45 (2.74%), and Pfs230 (1.84%), although low compared to those of pre-erythrocytic and erythrocytic stage antigens (Figure 6d; Supplementary Table 2).

Importantly, liver stage antigens ROM1, p36, and PALM had high epitope retention rates for both MHC I and MHC II, suggesting these proteins may be broadly immunogenic and can elicit both CD8+ and CD4+ T cell responses.

### Optimization of T cell epitope combinations for HLA and strain coverage

After initial epitope filtering, we used the TEpiNom tool to optimize the selection of MHC I and II epitopes for inclusion in combinations that maximized population coverage of HLA alleles. The algorithm prioritized epitopes that contributed the greatest marginal gains in HLA coverage while minimizing protein and domain redundancy.

For MHC I pre-erythrocytic stage epitopes, ten epitopes from HSP70-2, SLARP/SAP1, p36, FabZ, LISP1, LSA1, UIS3, p24_2, PL, and FabG achieved 98.15% coverage independently across HLA-A, HLA-B, and HLA-C alleles (Figure 7; Supplementary Table 3a). For MHC II epitopes, the number of epitopes needed to achieve the highest coverage varied across different life cycle stages (Figure 7). Pre-erythrocytic stage MHC II epitopes from PALM, AMA1, LISP1, TLP, CelTOS, and UIS3 collectively achieved 100% coverage across DRB1, DPA1-DPB1, and DQA1-DQB1 alleles (Figure 7; Supplementary Table 3b). In the erythrocytic stage, the maximum population coverage plateaued at 98.54% overall coverage with a combination of epitopes from RON2, VAR2CSA, PfSEA1, EBA-175, MSP3, PfRh5, Ripr, GLURP, and MSP1 (Figure 7; Supplementary Table 3c). Sexual stage MHC II epitope combinations, derived from Pfs230, along with Pfs48/45, Pfs47, and Pfs25, attained 97.84% overall coverage (Figure 7; Supplementary Table 3d).

**Figure 7.**
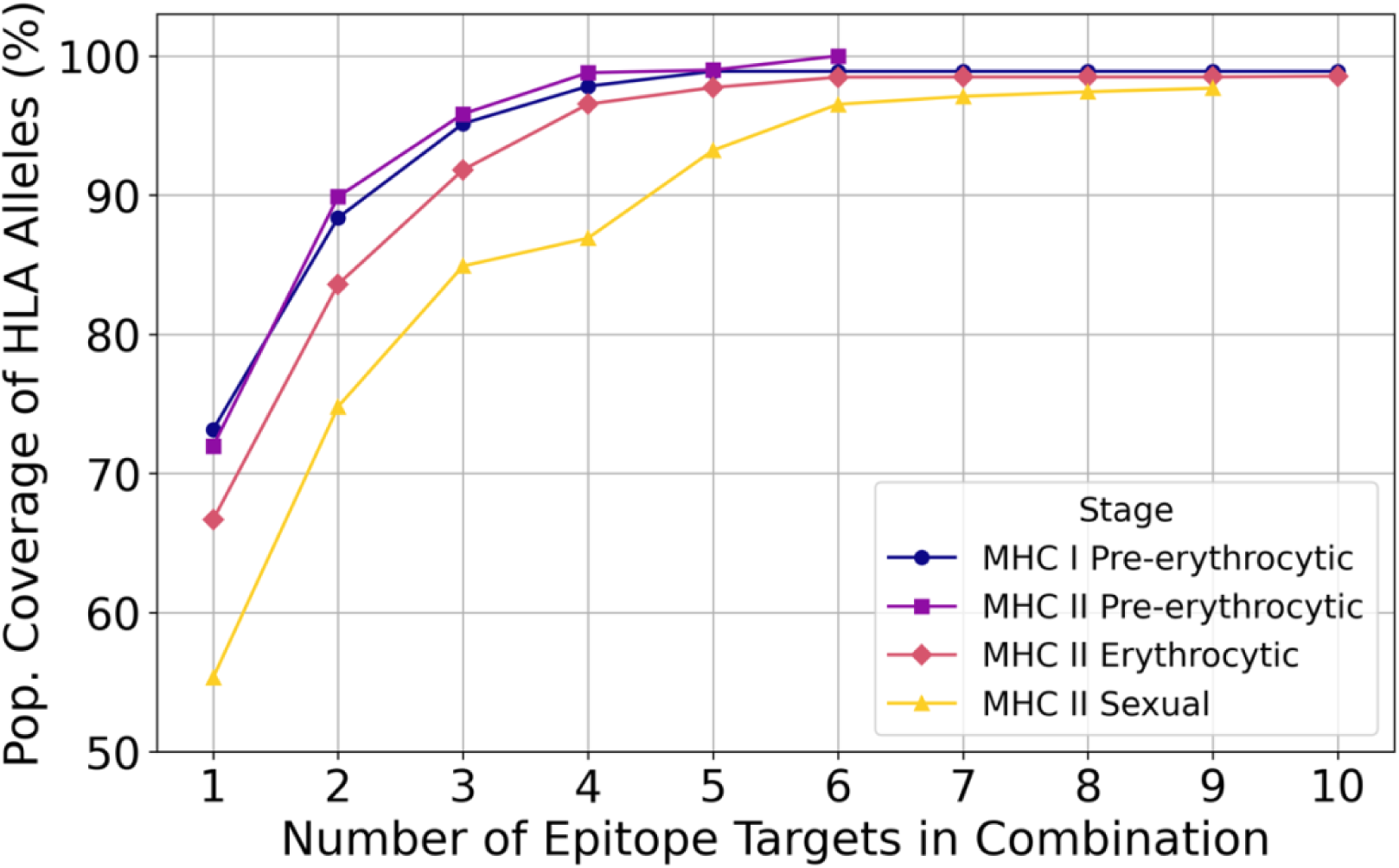
Population coverage of optimized epitope combinations. Population coverage (%) of HLA alleles for MHC I and MHC II epitope combinations of increasing number within each malaria life cycle stage. Epitope combinations were selected from filtered data sets based on conservation and binding affinity criteria. Each combination includes up to ten epitopes or stops if coverage reaches 100% or no longer increases.

### Association of HLA-Restricted T Cell Epitopes with Clinical Outcomes

To explore the relationship with HLA-restricted T cell epitopes and malaria clinical outcomes, we conducted a systematic literature review using various keyword combinations in PubMed searches to identify HLA alleles with significant associations with either protection or susceptibility to parasitemia, clinical malaria, or severe malaria outcomes. The search identified a total of 10 HLA alleles and nine allele groups associated with distinct clinical outcomes in malaria (Table 1).

**Table 1.**
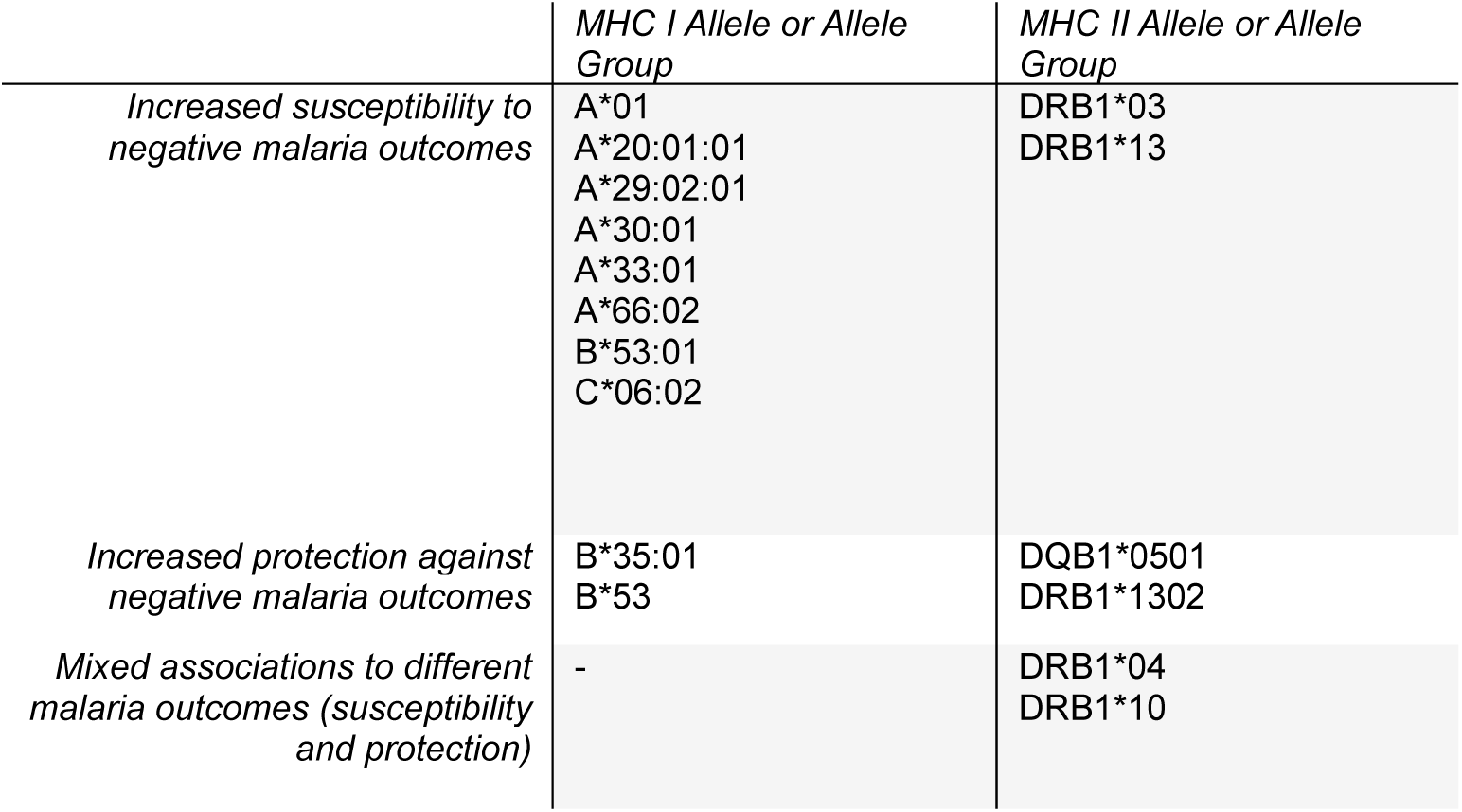
MHC I and II alleles and allele groups associated with malaria-related outcomes. HLA alleles with positive or negative associations to distinct malaria outcomes as found through a PubMed literature search are shown here along with if they were associated with increased susceptibility to, increased protection against, or a mixed association with different malaria outcomes.

More specifically, two MHC I alleles and four MHC I and II allele groups were associated with increased risk of either clinical malaria or parasitemia, while one MHC I allele and two MHC II allele groups were associated with protection against parasitemia. In the context of severe malaria in general or cerebral malaria specifically, 5 MHC I alleles and 2 MHC II allele groups were associated with increased risk, and one MHC I allele group and one MHC II allele were associated with protection (Supplementary Table 4). The MHC II allele HLA-DQB1*0501 was associated with decreased reinfection rates amongst children [25]. Predicted T cell epitopes that passed median binding affinity rank and epitope sequence conservation parameters were then analyzed for binding affinity to the identified HLA alleles and filtered to include epitopes predicted to bind strongly to non-protective and protective alleles. Amongst MHC I epitopes associated with susceptibility to parasitemia, uncomplicated malaria, or severe malaria, as predicted only in pre-erythrocytic stage proteins, no epitopes demonstrated promiscuous binding to all non-protective and protective alleles included in the analyses, and only one epitope was predicted to bind to all but one of the associated alleles. For MHC II epitopes that promiscuously bound to all non-protective and protective HLA alleles, 20 localized within pre-erythrocytic stage proteins, one within an erythrocytic stage protein, and four within sexual stage proteins. These results underscore predicted epitope binding restriction across HLA alleles associated with *P. falciparum* infection outcomes.

## Discussion

This study presents TEpiNom, a computational framework for rational T cell epitope selection in *Pf* vaccine development, integrating parasite antigen diversity and regional HLA allele frequencies to identify conserved, promiscuously binding epitopes with broad population coverage. Using this approach, we identified a set of MHC I and II epitopes across multiple parasite life stages with the potential to overcome antigenic diversity and HLA restriction, two major hurdles in malaria vaccine efficacy [5–7].

Our findings reinforce results from previous studies documenting HLA-restricted immune responses to *Pf* antigens and that a multi-epitope-based vaccine approach targeting conserved regions located across pre-erythrocytic, erythrocytic, and sexual stage proteins could generate a reliable and robust immune response in specific populations [7,15]. The current work adds to these earlier efforts that validated predicted CD4+ T cell epitopes in erythrocytic stage antigens, by broadening the scope of epitope prediction and down-selection to thousands of isolate sequences of 42 candidate malaria vaccine protein antigens against hundreds of endemic region HLA alleles [27]. Vaccine design informed by variant *Pf* sequences instead of the single laboratory strain upon which both RTS,S and R21 vaccines are based allows for identification of targets within circulating *Pf* strains causing disease. Integrating HLA allele frequencies derived from a wide range of sub-Saharan African populations ensures that the identified epitopes are immunologically relevant and optimally suited for populations most affected by malaria. This represents an innovative method aimed toward the rational design of epitope-based vaccines capable of eliciting robust immune responses in genetically diverse individuals, paving the road towards overcoming the hurdle of HLA restriction seen with RTS,S [7].

The potential for a multistage malaria vaccine is further reinforced by our findings on epitope conservation across multiple parasite life stages. This approach offers two distinct advantages. First, by targeting both pre-erythrocytic and erythrocytic stage antigens, a vaccine can protect against initial infection and progression to severe disease [45,46]. Second, including transmission-blocking epitopes derived from sexual stage proteins presents an opportunity to further reduce parasite circulation within the population, contributing to broader malaria eradication efforts at the community level [45,46]. In addition, the prioritization of epitopes that are non-overlapping in sequence and do not originate from the same protein antigen minimizes the risk of all targets included in a single vaccine becoming ineffective if the targeted protein antigen in a circulating parasite contains epitopes that do not match sequences in the vaccine construct. This comprehensive approach aligns with the broader goal of developing a malaria vaccine that is both protective at the individual level and capable of reducing community-wide transmission [46].

Existing T cell epitope tools can aid in T cell epitope down-selection; however, these tools independently focus on individual or select steps of the process [47–50]. IEDB’s TepiTool allows for the integration of T cell epitope prediction with downstream filtering for HLA allele binding affinity and epitope sequence conservation [47]. Another IEDB tool separately accomplishes population coverage analysis for a given set of epitopes and HLA alleles but does not complete any epitope combination optimization steps [48]. In contrast, TEpiNom can analyze large epitope data sets to down-select for conserved and broadly binding epitopes and then identify combinations of epitopes that maximize HLA allele coverage [49]. Technical University of Denmark’s PopCover 2.0 is a similar tool that incorporates epitope conservation and immune coverage [50] but may select for epitopes from the same antigen or that overlap in sequence, increasing the risk that parasite variation at a single antigen could evade vaccine-induced protection. Conversely, TEpiNom selects against combinations of epitopes originating from the same antigen or that overlap to promote a wide breadth of targeting in the final pool of vaccine candidates, a unique feature that reduces the risk of immune escape, which is common in complex pathogens such as *Pf*. Although a head-to-head comparison with existing tools would be necessary to evaluate differences in processing or efficiency, TEpiNom offers significant advantages through integrating epitope filtering parameters and prioritization steps, providing a more comprehensive and rational framework for epitope-based vaccine design.

Despite these advantages, limitations of the tools used and/or developed in this study exist. First, the number and identity of the protein antigens included in our analyses influence the down-selected epitope sets. Although we selected 42 well-described and high-priority *Pf* antigens across parasite life stages, this limited list adds an inherent constraint to the overall framework [33,34]. If key protective *Pf* antigens were excluded, promising epitopes may have been missed. Second, the down-selection process is dependent on the validity of epitope-HLA binding predictions made by computational tools, which, although incredibly advanced with high performance measures, may still introduce false positive or skewed predictions [31]. Third, HLA allele frequency inputs may not comprehensively reflect local population structures, especially in under-sampled regions [37], though HLA allele frequency data continues to be generated.

Additionally, when considering vaccine construct and deployment, practical constraints limit epitope-based vaccine design. While our tool nominates many conserved and immunogenic epitopes, only a limited number can feasibly be included in a single vaccine construct. Immunodominance and other considerations for a multi-epitope vaccine construct can be evaluated through experimental research. Examples of successful multivalent vaccines are the ones targeting *Streptococcus pneumoniae* that protect against more than 20 serotypes with a single injection [51].

From a pragmatic perspective, the selection of highly conserved and immunogenic epitopes with broad population coverage may reduce the current requirement for frequent booster doses, a significant advantage for vaccine distribution in malaria-endemic regions with limited public health resources. Additionally, our study highlights the need for careful consideration of HLA-restricted responses, as certain alleles may be associated with non-protective responses in the context of developing vaccine-mediated immunity, as demonstrated experimentally for CSP, in which immune responses to peptides were restricted amongst globally common HLA-DRB1 alleles [52]. By refining epitope selection based on predicted binding to protective HLA alleles, our findings contribute to the development of a highly promising next-generation malaria vaccine.

Beyond malaria vaccine development, the epitope down-selection tools presented in this study have broad applicability to a reverse vaccinology approach for other infectious diseases. The use of high-throughput and quantitative computational analyses to identify conserved and immunogenic epitopes can accelerate vaccine development for pathogens with high sequence variability, complex life cycles, or with high degrees of HLA-restricted immune responses, most notably HIV, dengue, and influenza [53–55]. Furthermore, the ability to rapidly prioritize epitopes based on sequence conservation and HLA binding promiscuity makes this approach highly relevant for pandemic preparedness. In the event of emerging infectious diseases, such tools could significantly accelerate vaccine development by identifying immunogenic targets with broad population coverage, facilitating streamlined vaccine development against novel pathogens. To realize this broader applicability of the TEpiNom tool, a key next step is expanding and standardizing the tool and overall workflow to facilitate pathogen-specific input data while maintaining updated, region-specific HLA allele frequency datasets.

This study demonstrates the feasibility of using large-scale computational approaches to guide next-generation malaria vaccine development. By integrating parasite sequence diversity data, endemic area HLA allele frequencies, and epitope presentation predictions, we have designed a strategy that enhances the likelihood of developing an effective, scalable, and widely applicable vaccine that could provide durable protection against diverse *Pf* variants in endemic settings. Advancing computational predictions requires laboratory validation to confirm the immunogenicity of prioritized epitopes via MHC stabilization assays, T cell stimulation studies, and eventually in a more biologically relevant context using *in vivo* testing in preclinical models to assess for immunogenic and protective potential, similar to previous studies that validated *Pf* epitopes [32,56–58].

In parallel, further refinement of the epitope prioritization criteria should continue based on results of *in vitro* and *ex vivo* analyses, including those that are publicly available in databases such as the Immune Epitope Database (IEDB), as these experiments can inform the rational adjustment of weighting or prioritizing certain criteria based on real-world relevance [59]. This study provides a critical step toward the rational design of malaria vaccines that are both broadly protective and capable of addressing the complex challenges posed by *Pf* infection.

## Acknowledgements

This publication was made possible by the University of Maryland Baltimore Institute for Clinical and Translational Research (ICTR) which is funded in part by Grant Number T32 TR004928 from the National Center for Advancing Translational Sciences (NCATS), a component of the National Institutes of Health (NIH), and NIH Roadmap for Medical Research. Its contents are solely the responsibility of the authors and do not necessarily represent the official view of the University of Maryland, Baltimore ICTR, NCATS, or NIH. MBL is supported by grants and contracts to his institution from the U.S. National Institutes of Health (UM1AI148689 and U01AI155300), Bill & Melinda Gates Foundation (INV-030857), Bill & Melinda Gates Medical Research Institute, and BioNTech, SE. The funders had no role in study design, data collection and analysis, decision to publish, or preparation of the manuscript.

## Data availability statement

Generated protein sequence and HLA allele data sets are available upon request. The T Cell Epitope Nomination (TEpiNom; Patent Pending) tool for down-selection and optimization is available at https://github.com/alexlaurenson/epiweight.

---

**Supplementary Table 1.**
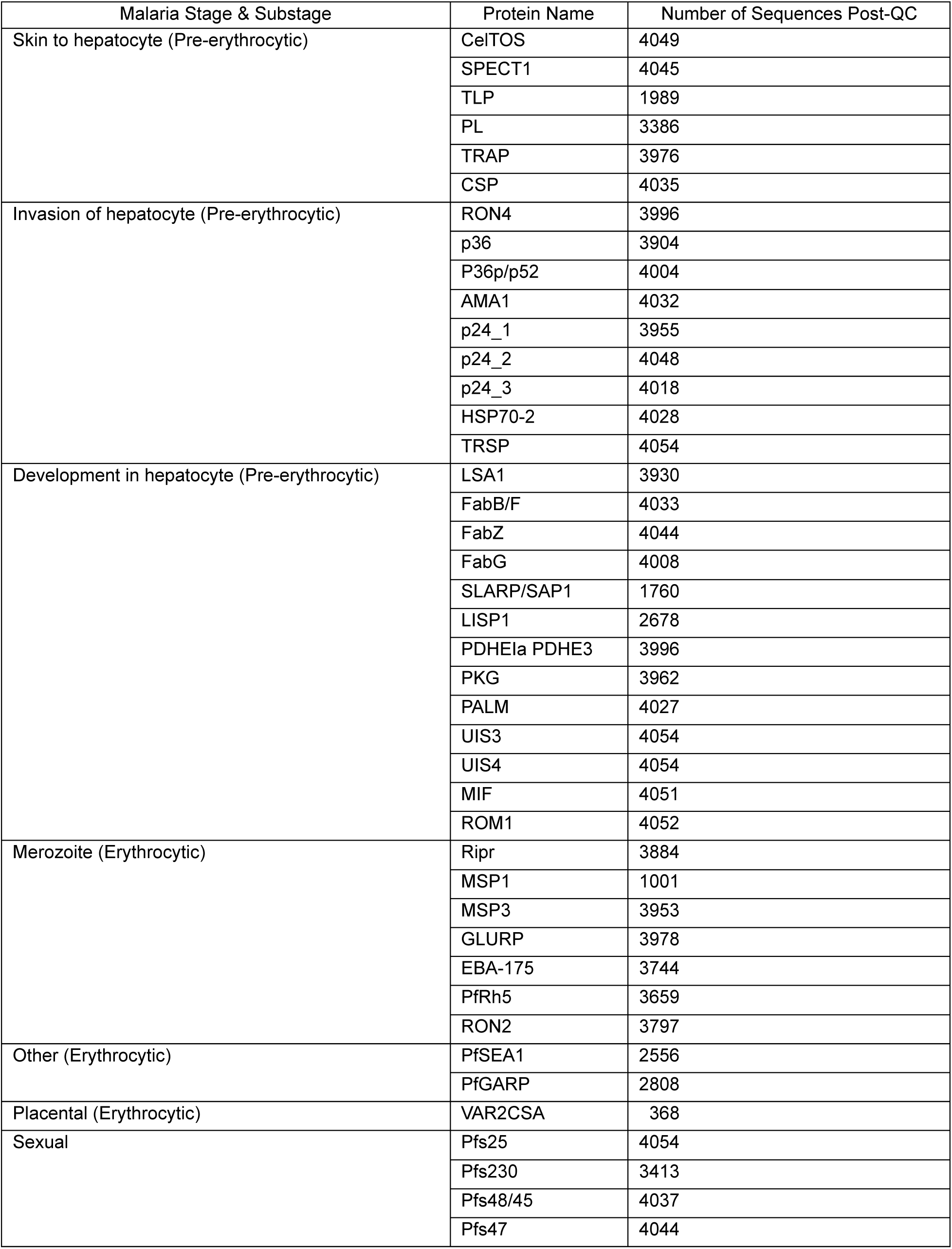
Protein sequence data set per malaria antigen. Number of sample isolates in each protein’s sequence data set after quality control filtering. Sequences were acquired from MalariaGEN *Plasmodium falciparum* genomic data set and filtered out if within-infection fixation index (F_WS_) was greater than 0.95 indicating a potential polyclonal infection or if the resulting consensus sequence contained biologically improbable nonsense mutations.

**Supplementary Table 2.**
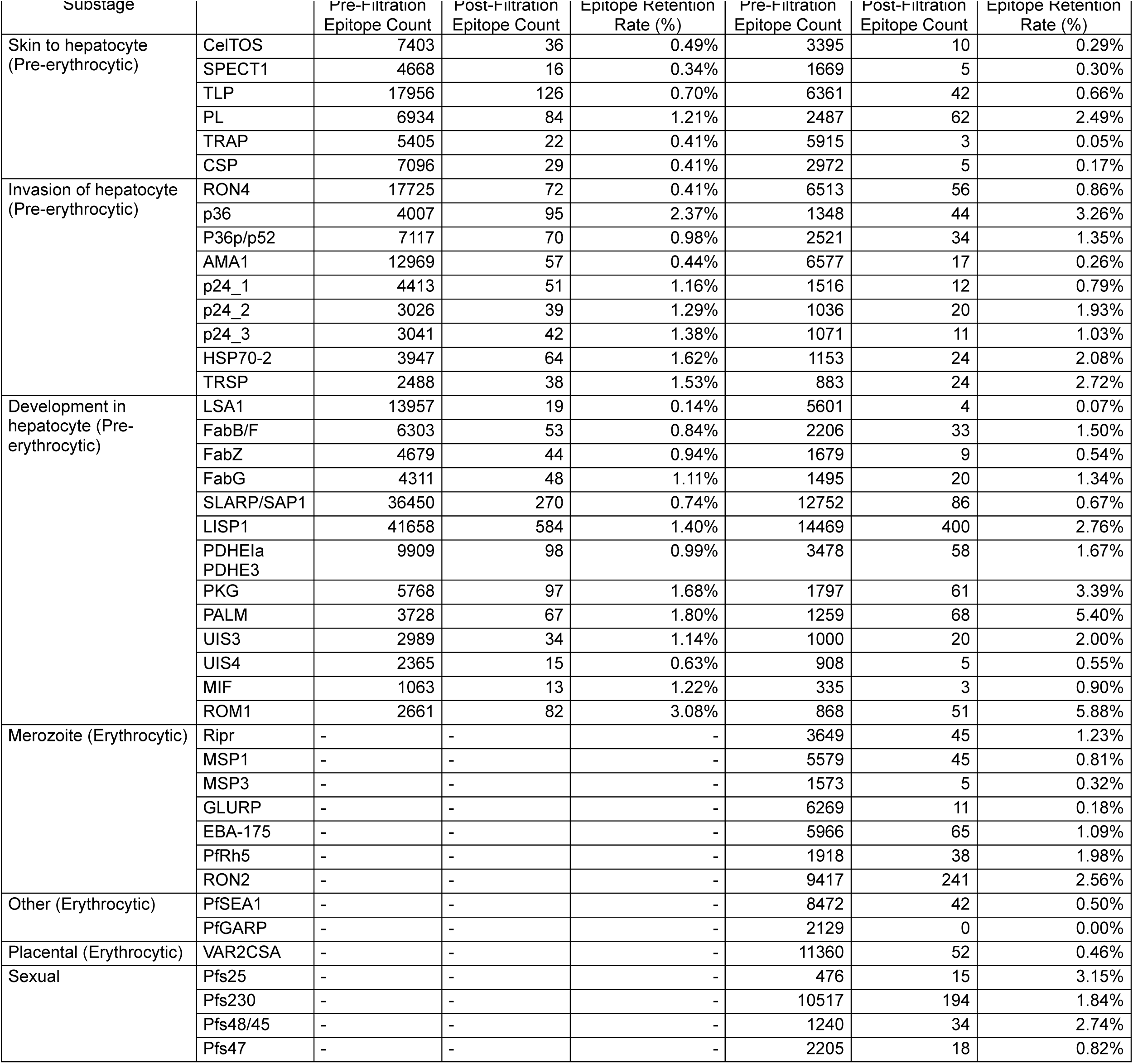
Predicted and retained T cell epitopes across malaria vaccine candidate proteins. Number of predicted MHC I and MHC II epitopes before and after applying conservation and binding affinity filters (<10% median binding rank, >95% conservation). Data is categorized by malaria life cycle stage and substage with epitope counts shown for each protein after filtration steps. The percentage of retained epitopes after filtration is also provided for both MHC I and MHC II predictions as an epitope retention rate. MHC I epitope predictions performed exclusively for pre-erythrocytic stage proteins.

**Supplementary Table 3.**
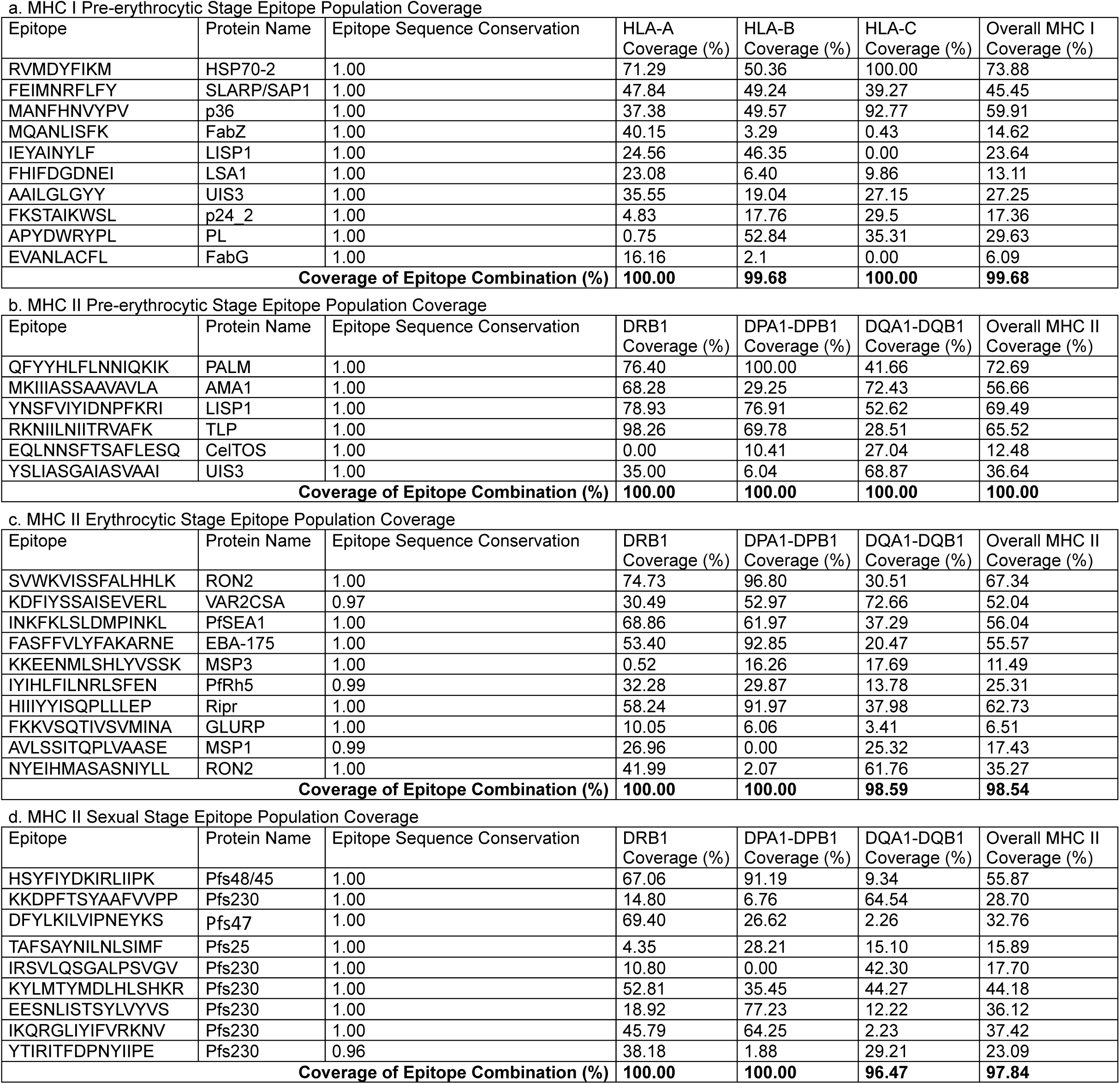
Epitope combinations for optimized MHC population coverage. MHC I or MHC II epitope combinations within each malaria life cycle stage found to maximize coverage of HLA alleles within endemic region population selecting from epitope data set after median binding affinity rank and epitope sequence conservation filtration steps. Epitope-HLA population coverage (%) shown for each relevant HLA class within MHC I or II with Coverage of Epitope Combination (%) calculated by summing unique HLA coverage of given epitope combination.

**Supplementary Table 4.**
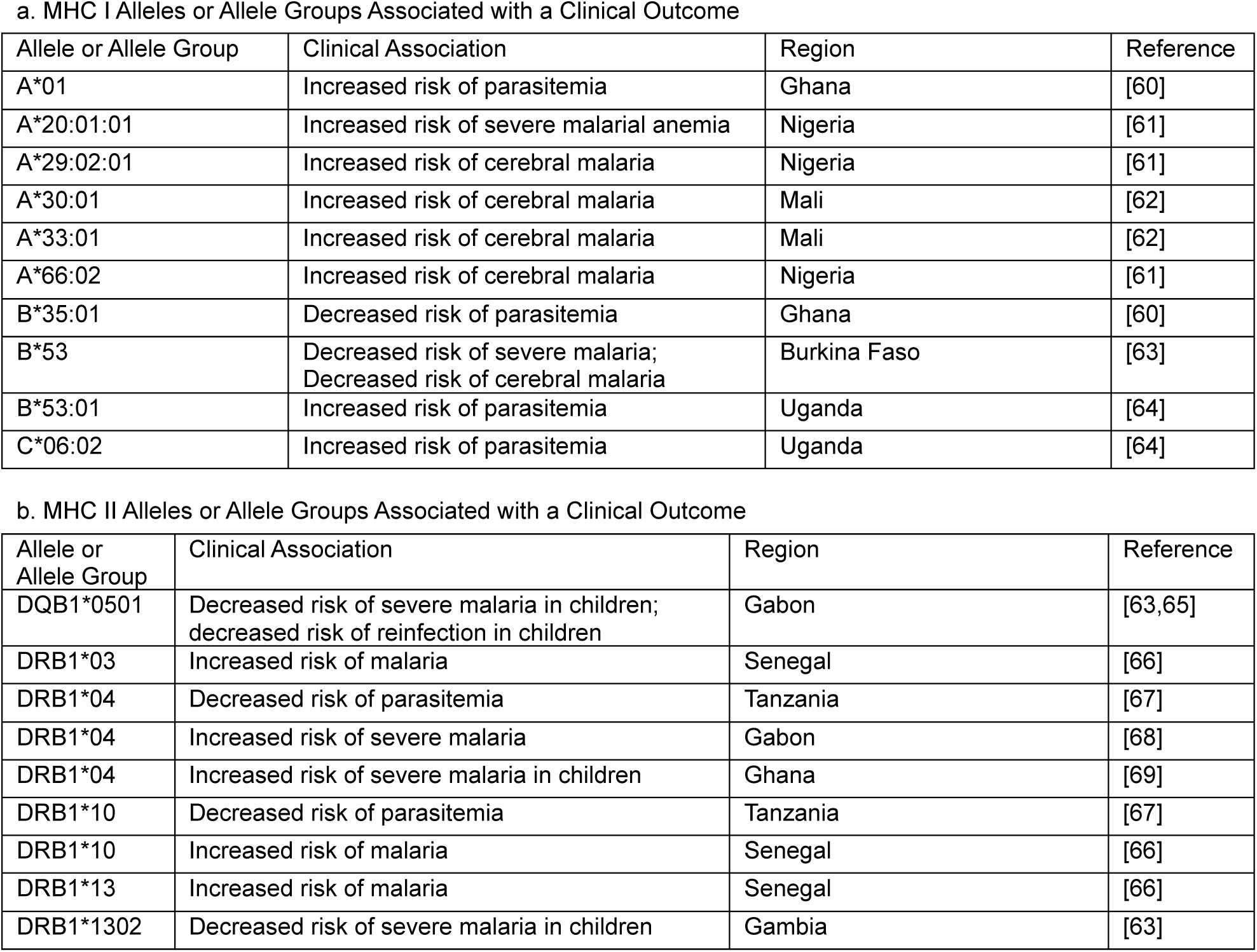
MHC I and II alleles and allele groups associated with clinical outcomes. HLA alleles with positive or negative associations to distinct malaria clinical outcomes as found through a PubMed literature search are shown here along with specific outcome, country from which the data originated, and literature reference.

## Notes

https://github.com/alexlaurenson/epiweight

### Competing Interest Statement

The authors have declared no competing interest.

## References

1. WHO. World malaria report 2024. (2024).

2. Moorthy, V., Hamel, M. J. & Smith, P. G. Malaria vaccines for children: and now there are two. The Lancet 403, 504–505 (2024).

3. Olotu, A. et al. Seven-Year Efficacy of RTS,S/AS01 Malaria Vaccine among Young African Children. N. Engl. J. Med. 374, 2519–2529 (2016).

4. Datoo, M. S. et al. Efficacy of a low-dose candidate malaria vaccine, R21 in adjuvant Matrix-M, with seasonal administration to children in Burkina Faso: a randomised controlled trial. The Lancet 397, 1809–1818 (2021).

5. Takala, S. L. & Plowe, C. V. Genetic diversity and malaria vaccine design, testing and efficacy: preventing and overcoming ‘vaccine resistant malaria’. Parasite Immunol. 31, 560– 573 (2009).

6. Ouattara, A. et al. Designing malaria vaccines to circumvent antigen variability. Vaccine 33, 7506–7512 (2015).

7. Nielsen, C. M. et al. RTS,S malaria vaccine efficacy and immunogenicity during Plasmodium falciparum challenge is associated with HLA genotype. Vaccine 36, 1637–1642 (2018).

8. Draper, S. J. et al. Malaria Vaccines: Recent Advances and New Horizons. Cell Host Microbe 24, 43–56 (2018).

9. Miura, K. Progress and prospects for blood-stage malaria vaccines. Expert Rev. Vaccines 15, 765–781 (2016).

10. Gardner, M. J. et al. Genome sequence of the human malaria parasite Plasmodium falciparum. Nature 419, 498–511 (2002).

11. Dame, J. B. et al. Current status of the Plasmodium falciparum genome project. Mol. Biochem. Parasitol. 79, 1–12 (1996).

12. Shah, Z. et al. Whole-genome analysis of Malawian Plasmodium falciparum isolates identifies possible targets of allele-specific immunity to clinical malaria. PLOS Genet. 17, e1009576 (2021).

13. Takala, S. L. et al. Extreme Polymorphism in a Vaccine Antigen and Risk of Clinical Malaria: Implications for Vaccine Development. Sci. Transl. Med. 1, (2009).

14. MalariaGEN et al. Pf7: an open dataset of Plasmodium falciparum genome variation in 20,000 worldwide samples. Wellcome Open Res. 8, 22 (2023).

15. Thera, M. A. et al. A Field Trial to Assess a Blood-Stage Malaria Vaccine. N. Engl. J. Med. 365, 1004–1013 (2011).

16. Ouattara, A. et al. Molecular Basis of Allele-Specific Efficacy of a Blood-Stage Malaria Vaccine: Vaccine Development Implications. J. Infect. Dis. 207, 511–519 (2013).

17. Bailey, J. A. et al. Microarray analyses reveal strain-specific antibody responses to Plasmodium falciparum apical membrane antigen 1 variants following natural infection and vaccination. Sci. Rep. 10, 3952 (2020).

18. Genton, B. et al. A Recombinant Blood-Stage Malaria Vaccine Reduces *Plasmodium falciparum* Density and Exerts Selective Pressure on Parasite Populations in a Phase 1–2b Trial in Papua New Guinea. J. Infect. Dis. 185, 820–827 (2002).

19. Rappuoli, R., Bottomley, M. J., D’Oro, U., Finco, O. & De Gregorio, E. Reverse vaccinology 2.0: Human immunology instructs vaccine antigen design. J. Exp. Med. 213, 469–481 (2016).

20. Soria-Guerra, R. E., Nieto-Gomez, R., Govea-Alonso, D. O. & Rosales-Mendoza, S. An overview of bioinformatics tools for epitope prediction: implications on vaccine development. J. Biomed. Inform. 53, 405–414 (2015).

21. Rosendahl Huber, S., Van Beek, J., De Jonge, J., Luytjes, W. & Van Baarle, D. T Cell Responses to Viral Infections - Opportunities for Peptide Vaccination. Front. Immunol. 5, (2014).

22. Kimura, K. et al. CD8^+^ T Cells Specific for a Malaria Cytoplasmic Antigen Form Clusters around Infected Hepatocytes and Are Protective at the Liver Stage of Infection. Infect. Immun. 81, 3825–3834 (2013).

23. Cockburn, I. A. et al. In vivo imaging of CD8^+^ T cell-mediated elimination of malaria liver stages. Proc. Natl. Acad. Sci. 110, 9090–9095 (2013).

24. Su, Z. & Stevenson, M. M. Central Role of Endogenous Gamma Interferon in Protective Immunity against Blood-Stage *Plasmodium chabaudi* AS Infection. Infect. Immun. 68, 4399– 4406 (2000).

25. Horowitz, A. et al. Cross-Talk between T Cells and NK Cells Generates Rapid Effector Responses to *Plasmodium falciparum -* Infected Erythrocytes. J. Immunol. 184, 6043–6052 (2010).

26. Oakley, M. S. et al. T-bet modulates the antibody response and immune protection during murine malaria. Eur. J. Immunol. 44, 2680–2691 (2014).

27. Pérez-Mazliah, D. et al. Follicular Helper T Cells are Essential for the Elimination of Plasmodium Infection. EBioMedicine 24, 216–230 (2017).

28. Pérez-Mazliah, D. et al. Disruption of IL-21 Signaling Affects T Cell-B Cell Interactions and Abrogates Protective Humoral Immunity to Malaria. PLOS Pathog. 11, e1004715 (2015).

29. Hughes, A. L. & Nei, M. Pattern of nucleotide substitution at major histocompatibility complex class I loci reveals overdominant selection. Nature 335, 167–170 (1988).

30. Hughes, A. L. & Nei, M. Nucleotide substitution at major histocompatibility complex class II loci: evidence for overdominant selection. Proc. Natl. Acad. Sci. 86, 958–962 (1989).

31. Reynisson, B., Alvarez, B., Paul, S., Peters, B. & Nielsen, M. NetMHCpan-4.1 and NetMHCIIpan-4.0: improved predictions of MHC antigen presentation by concurrent motif deconvolution and integration of MS MHC eluted ligand data. Nucleic Acids Res. 48, W449– W454 (2020).

32. Kotraiah, V. et al. Identification and Immune Assessment of T Cell Epitopes in Five Plasmodium falciparum Blood Stage Antigens to Facilitate Vaccine Candidate Selection and Optimization. Front. Immunol. 12, 690348 (2021).

33. Duffy, P. E., Sahu, T., Akue, A., Milman, N. & Anderson, C. Pre-erythrocytic malaria vaccines: identifying the targets. Expert Rev. Vaccines 11, 1261–1280 (2012).

34. Duffy, P. E. & Patrick Gorres, J. Malaria vaccines since 2000: progress, priorities, products. Npj Vaccines 5, 48 (2020).

35. Danecek, P. et al. Twelve years of SAMtools and BCFtools. GigaScience 10, giab008 (2021).

36. Perez, G. et al. The UCSC Genome Browser database: 2025 update. Nucleic Acids Res. gkae974 (2024). doi:10.1093/nar/gkae974

37. Gonzalez-Galarza, F. F. et al. Allele frequency net database (AFND) 2020 update: gold-standard data classification, open access genotype data and new query tools. Nucleic Acids Res. gkz1029 (2019). doi:10.1093/nar/gkz1029

38. Modiano, D. et al. HLA class I in three West African ethnic groups: genetic distances from sub-Saharan and Caucasoid populations: HLA class I in African ethnic groups. Tissue Antigens 57, 128–137 (2001).

39. Kijak, G. H. et al. HLA class I allele and haplotype diversity in Ugandans supports the presence of a major east African genetic cluster. Tissue Antigens 73, 262–269 (2009).

40. Katoh, K. & Standley, D. M. MAFFT multiple sequence alignment software version 7: improvements in performance and usability. Mol. Biol. Evol. 30, 772–780 (2013).

41. Fu, L., Niu, B., Zhu, Z., Wu, S. & Li, W. CD-HIT: accelerated for clustering the next-generation sequencing data. Bioinformatics 28, 3150–3152 (2012).

42. Rossum, G. van & Drake, F. L. *The Python language reference*. (Python Software Foundation, 2010).

43. The pandas development team. pandas-dev/pandas: Pandas. (2024). doi: 10.5281/ZENODO.3509134

44. Harris, C. R. et al. Array programming with NumPy. Nature 585, 357–362 (2020).

45. Ouattara, A. & Laurens, M. B. Vaccines Against Malaria. Clin. Infect. Dis. 60, 930–936 (2015).

46. Malaria Vaccines: Preferred Product Characteristics and Clinical Development Considerations. (World Health Organization, 2022).

47. Paul, S., Sidney, J., Sette, A. & Peters, B. TepiTool: A Pipeline for Computational Prediction of T Cell Epitope Candidates. Curr. Protoc. Immunol. 114, (2016).

48. Bui, H.-H. et al. Predicting population coverage of T-cell epitope-based diagnostics and vaccines. BMC Bioinformatics 7, 153 (2006).

49. Bui, H.-H., Sidney, J., Li, W., Fusseder, N. & Sette, A. Development of an epitope conservancy analysis tool to facilitate the design of epitope-based diagnostics and vaccines. BMC Bioinformatics 8, 361 (2007).

50. Nilsson, J. B., Grifoni, A., Tarke, A., Sette, A. & Nielsen, M. PopCover-2.0. Improved Selection of Peptide Sets With Optimal HLA and Pathogen Diversity Coverage. Front. Immunol. 12, 728936 (2021).

51. Nielsen, K. F. et al. Follow-Up Study of Effectiveness of 23-Valent Pneumococcal Polysaccharide Vaccine Against All-Type and Serotype-Specific Invasive Pneumococcal Disease, Denmark. Emerg. Infect. Dis. 30, (2024).

52. Doolan, D. L. et al. HLA-DR-Promiscuous T Cell Epitopes from *Plasmodium falciparum* Pre-Erythrocytic-Stage Antigens Restricted by Multiple HLA Class II Alleles. J. Immunol. 165, 1123–1137 (2000).

53. Bronke, C. et al. HIV escape mutations occur preferentially at HLA-binding sites of CD8 T-cell epitopes. AIDS 27, 899–905 (2013).

54. Loke, H. et al. Strong HLA Class I–Restricted T Cell Responses in Dengue Hemorrhagic Fever: A Double-Edged Sword? J. Infect. Dis. 184, 1369–1373 (2001).

55. Hensen, L. et al. CD8+ T cell landscape in Indigenous and non-Indigenous people restricted by influenza mortality-associated HLA-A*24:02 allomorph. Nat. Commun. 12, 2931 (2021).

56. Bergmann-Leitner, E. S. et al. Computational and Experimental Validation of B and T-Cell Epitopes of the In Vivo Immune Response to a Novel Malarial Antigen. PLoS ONE 8, e71610 (2013).

57. Quadiri, A., Kalia, I., Kashif, M. & Singh, A. P. Identification and characterization of protective CD8^+^ T-epitopes in a malaria vaccine candidate SLTRiP. Immun. Inflamm. Dis. 8, 50–61 (2020).

58. Ganeshan, H. et al. Measurement of ex vivo ELISpot interferon-gamma recall responses to Plasmodium falciparum AMA1 and CSP in Ghanaian adults with natural exposure to malaria. Malar. J. 15, 55 (2016).

59. Vita, R. et al. The Immune Epitope Database (IEDB): 2024 update. Nucleic Acids Res. 53, D436–D443 (2025).

60. Yamazaki, A. et al. Human leukocyte antigen class I polymorphisms influence the mild clinical manifestation of Plasmodium falciparum infection in Ghanaian children. Hum. Immunol. 72, 881–888 (2011).

61. Ademola, S. A., Amodu, O. K. & Yindom, L. HLA-A Alleles Differentially Associate with Severity to Plasmodium falciparum Malaria Infection in Ibadan, Nigeria. Afr. J. Biomed. Res. 20, 223–228 (2017).

62. Lyke, K. E. et al. Association of HLA alleles with *Plasmodium falciparum* severity in Malian children. Tissue Antigens 77, 562–571 (2011).

63. Hill, A. V. S. et al. Common West African HLA antigens are associated with protection from severe malaria. Nature 352, 595–600 (1991).

64. Digitale, J. C. et al. HLA Alleles B*53:01 and C*06:02 Are Associated With Higher Risk of P. falciparum Parasitemia in a Cohort in Uganda. Front. Immunol. 12, 650028 (2021).

65. May, J., Lell, B., Luty, A. J. F., Meyer, C. G. & Kremsner, P. G. HLA-DQB1*0501–Restricted Th1 Type Immune Responses to *Plasmodium falciparum* Liver Stage Antigen 1 Protect against Malaria Anemia and Reinfections. J. Infect. Dis. 183, 168–172 (2001).

66. Ndiaye, M. et al. Susceptibility to neuro-malaria and HLA-DR alleles in Senegal. Dakar Med. 43, 25–28 (1998).

67. Bennett, S. et al. Human leucocyte antigen (HLA) and malaria morbidity in a Gambian community. Trans. R. Soc. Trop. Med. Hyg. 87, 286–287 (1993).

68. May, J. et al. HLA Class II Factors Associated with *Plasmodium falciparum* Merozoite Surface Antigen Allele Families. J. Infect. Dis. 179, 1042–1045 (1999).

69. Osafo-Addo, A. D. et al. HLA-DRB1*04 allele is associated with severe malaria in northern Ghana. Am. J. Trop. Med. Hyg. 78, 251–255 (2008).

